# Digitalized organoids: integrated pipeline for 3D high-speed analysis of organoid structures using multilevel segmentation and cellular topology

**DOI:** 10.1101/2023.11.08.566158

**Authors:** Hui Ting Ong, Esra Karatas, Gianluca Grenci, Florian Dilasser, Saburnisha Binte Mohamad Raffi, Damien Blanc, Titouan Poquillon, Elise Drimaracci, Dimitri Mikec, Cora Thiel, Oliver Ullrich, Victor Racine, Anne Beghin

**Author notes:** co-first authors (Hui Ting Ong, Esra Karatas). **Corresponding author:** Anne Beghin.

## Abstract

Analysing the tissue morphogenesis and function is crucial for unravelling the underlying mechanisms of tissue development and disease. Organoids, 3D *in vitro* models that mimic the architecture and function of human tissues, offer a unique opportunity to study effects of external perturbators that are difficult to replicate *in vivo*. However, large-scale screening procedures for studying the effects of different ‘stress’ on cellular morphology and topology of these 3D tissue-like system face significant challenges, including limitations in high-resolution 3D imaging, and accessible 3D analysis platforms. These limitations impede the scale and throughput necessary to accurately quantify the effects of mechanical and chemical cues. Here, we present a novel, fine-tuned pipeline for screening morphology and topology modifications in 3D cell culture using multilevel segmentations and cellular topology, based on confocal microscopy and validated across different image qualities. Our pipeline incorporates advanced image analysis algorithms and artificial intelligence (AI) for multiscale 3D segmentation, enabling quantification of morphology changes at both the nuclear and cytoplasmic levels, as well as at the organoid scale. Additionally, we investigate cell relative position and employ neighbouring topology analysis to identify tissue patterning and their correlations with organoid microniches. Eventually, we have organized all the extracted features, 3D segmented masks and raw images into a single database to allow statistical and data mining approaches to facilitate data analysis, in a biologist-friendly way. We validate our approach through proof-of-concept experiments, including well-characterized conditions and poorly explored mechanical stressors such as microgravity, showcasing the versatility of our pipeline. By providing a powerful tool for discovery-like assays in screening 3D organoid models, our pipeline has wide-ranging interests from biomedical applications in development and aging-related pathologies to tissue engineering and regenerative medicine.

## INTRODUCTION

The capability to accurately measure three-dimensional (3D) deformations of cells within their native 3D structures or tissues is of paramount significance for studying within the realms of tissue’s health, dynamics, and pathological conditions. While normative mechanical forces and chemical cues are essential for the orchestration of the tissue maturation and optimal functionality, the advent of aberrant stresses augments the propensity occurrence of deleterious consequences. Research has firmly established that these stresses can accelerate tissue aging, induce pathophysiological changes, and hinder tissue regeneration^1,2^.

In this context, organoids have emerged as a crucial instrument of choice. Organoids characterized as autonomous, three-dimensional cellular assemblies emulating the architectural and functional nuances of specific basic blocks of organs have been crucial to decipher modifications of genetic expression, cell communication pathway, and the provocation of cellular damage in response to external cues. By furnishing an ethically and methodologically viable platform, they facilitate the investigation of diverse signalling effects on tissues, enabling precise external stimulations that are operationally intricate within animal models^3–5^. A marked departure from conventional two-dimensional cell cultures, these dynamic three-dimensional structures reverberate with a heightened physiological relevance^6,7^. Notably, the precise definition of organoid geometry, fundamental in shaping patterning and cryptogenesis, could be achieved through bioengineered extracellular microniches of meticulous geometries and stiffness^8^. While studies have demonstrated external microniche and mechanical stimulation’s feasibility, they primarily focus on overall morphological descriptors of whole organoid, or 2D descriptors and gene expression, and are lacking cellular 3D morphology quantification^9–11^.

Crucially, the cornerstone of describing cell (and/or intracellular compartments) deformation relies on precise image-based segmentation techniques. This capacity to gauge cellular deformations must derive from the utilization of ubiquitous cellular markers present across all cell types, thus ensuring maximum independence from cellular differentiation and phenotypic variations. The analysis of complex morphological changes at different scales (single cells, clusters of cells and whole organoids) in combination with approaches based on high content 3D imaging and screening methods has the potential to reveal new mechanisms of response or adaptation during tissue development, injuries and aging related diseases.

In the domain of biomedical research and screening campaigns, the demand for accurate 3D cell segmentation from microscopic images has led to the development of various open-source deep learning-based solutions. Various software for these purposes have been developed in recent years, including Ilastik^12^ and OrganoidTracker (nuclei only)^13^. Most of these have Fiji or CellProfiler opensource plug-ins, which make it more accessible to the scientific community. These softwares usually contain functions for organoids analysis such as image segmentation and object classification based on phenotypic markers. However, existing solutions require high-quality three-dimensional image resolution for robust and accurate segmentation and the training phase for these solutions is often time-consuming and does not readily accommodate changes in resolution, staining, cell types or specific image characteristics. Notably for the cell contour, the recent introduction of 3DCellSeg, for segmentation of vegetal or zebrafish 3D structures, has garnered attention for its robustness and precision compared to other existing methods^14^. However, it is worth noting that the training and evaluation of 3DCellSeg and others available deep learning algorithms has primarily been conducted on image datasets that significantly differ from the challenging real-world conditions encountered in biomedical research, where resolutions and signal-to-noise ratios are considerably lower, particularly in the case of mammalian cell organoids. These limitations currently impede the applicability of existing deep learning method for precise segmentation of compact organoid cells in practical scenarios, with the added concern that authors habitually did not disclose computational time requirements for the segmentation process.

In this context, we introduce an innovative, comprehensive methodology designed to assess alterations in both morphology and topology of cells within 3D organoids. This approach integrates multilevel segmentation and cellular topology techniques, leveraging common confocal microscope imagery compatible with screening assay and validated across various image qualities. Our pipeline incorporates sophisticated image analysis algorithms for multiscale 3D segmentation together with a database architecture facilitating the quantification of morphological changes at nuclear, cytoplasmic, and organoid levels. We based our approach on universal markers (DNA and Actin staining) to ensure independence from specific phenotype profiles and enable researchers to avoid the necessity of utilizing immunostaining long procedure for these morphological features. Furthermore, our exploration extends to cell positional relationships, encompassing tissue patterning detection to unveil intricate morphological attributes. Our proposed solution enables the generation of numerous morphological and topological descriptors while concurrently allowing the reduction of the dimensionality of this database to generate complex signatures relevant to conditions devoid of prior assumptions (i.e screening or discovery-like assay). These signatures can be biologically interpreted and readily validated through a biologist-friendly manipulation pipeline.

Our method enables streamlined analysis through multilevel 3D segmentation of numerous organoids incorporating descriptors of internal Cell-to-Cell and Cell-to-Neighbourhood organization and statistical extraction suites, thus enabling the unlocking of organoids’ potential to fulfil their complete roles in drug screening, understanding diseases, tissue aging and regeneration. There are 3 innovated parts in this pipeline which consists of (i) a fast-and-reliable 3D segmentation adapted to the real-lab world conditions, (ii) 3D topology descriptors (Cell-to-Neighbourhood) analysis to quantitatively identify tissue patterning and, (iii) generation of comprehensive 3D cell morphological signatures allowing the assessment of different conditions of mechanical constraints.

## RESULTS

### 1/ Concept and benchmarking

We describe the principle of “digitalized” organoids, showcasing the capabilities of our analysis techniques (**Figure 1**). Our approach is tailored to commonly used 3D fluorescence microscopy methods, such as spinning disk or point scanner confocal and ensuring compatibility with large batches of organoids (several hundreds). Notably, we have restricted the image acquisition process to be completed in less than a minute per organoid for 2-color images, often resulting in a larger z-step (e.g. 3 µm). This restricted time establishes our approach as a benchmark for high-throughput screening of organoids.

**Figure 1:**
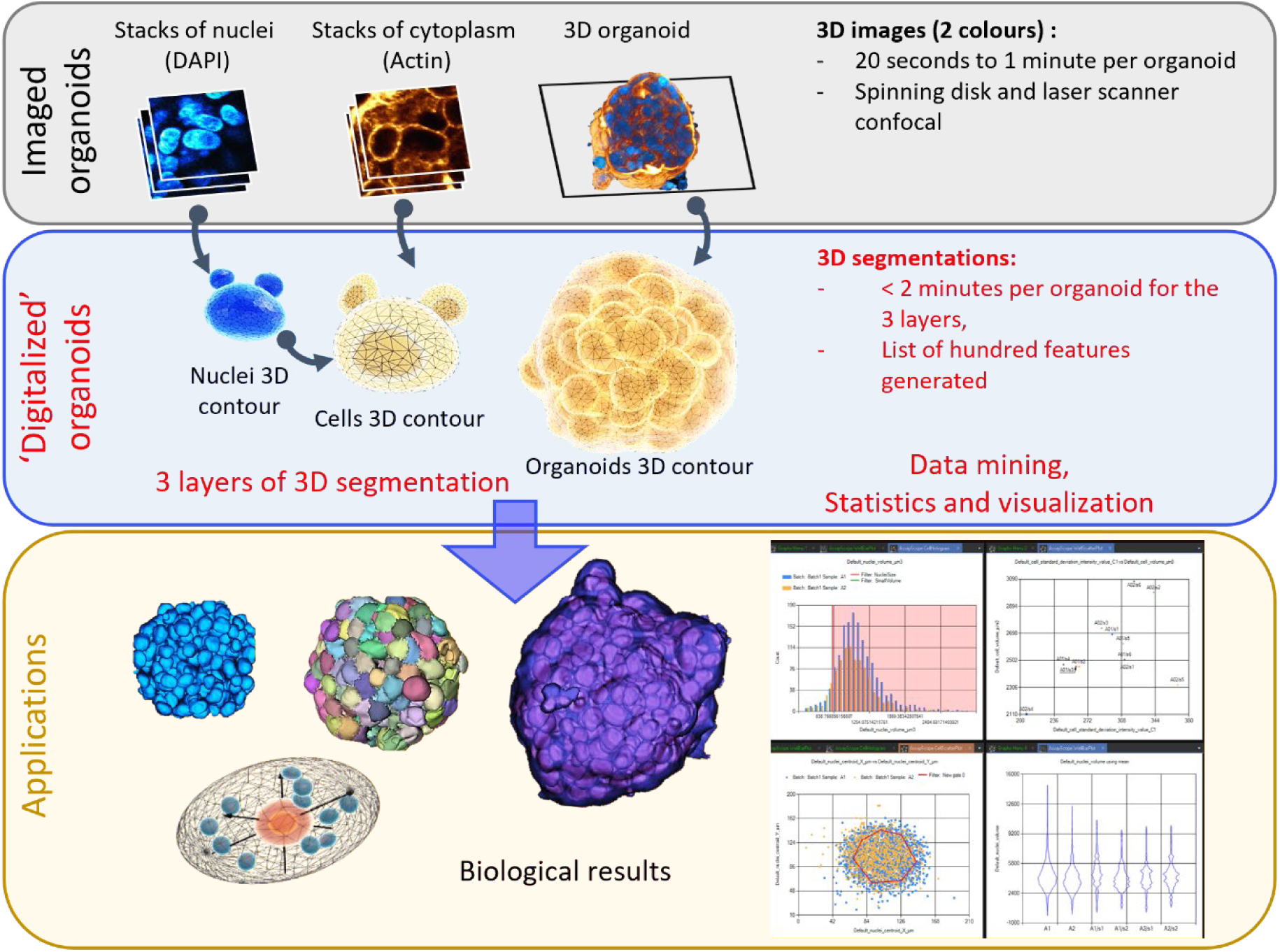
’Digitalized’ organoids principle. Top panel: Organoids were imaged using state-of-the-art 3D fluorescence microscopy techniques compatible with screening approaches (such as spinning disk or laser scanner confocal) at a speed of less than a minute per organoid for complete multicolor 3D stacks. **Middle panel:** The complete 3D digitalization of organoids consists of three layers of individual cell segmentation: nuclei contours from DAPI images (AI-based analysis), cell contours using both the nuclei contours and the raw images of actin, and the whole organoid contour obtained from the raw channels. Altogether, these 3D segmentations are performed in less than 2 minutes per organoid, without the need for any GPU-based processes, and generate a list of hundreds of morphological and topological descriptors. **Low panel:** The resulting database, the 3D segmentation layers, and the raw images have been made understandable through a dedicated data mining software (cytometry-like software) that enables the extraction of biological results and the feedback on raw images.

The 3D digitalization, from sub-cellular level to whole organoid scale, involves three levels of segmentation and has been compiled by using open-source software and available AI networks. Firstly, nuclei surfaces are extracted using AI-based analysis of DAPI images, which is based on a finely trained 3D Stardist CNN^15^ model using simulated images that closely mimic real-world conditions. We have chosen Stardist as it represents a state-of-the-art approach for nuclei segmentation well known for its accuracy and speed. Among other 3D instance segmentation frameworks such as U-Net3d^16^, Cellpose3d^17^, EmbedSeg^18^, Stardist has proven to perform extremely well for convex object such as nuclei, especially in complex biological context with dense group of nuclei and noisy background, while being comparatively faster than its counterparts^18,19^. However, it does have limitations, particularly when applied to 3D segmentation. Our fine-tuned model has been carefully designed to tolerate variations in intensities, contrast, and sizes (**Online Methods**). Secondly, cell surfaces are obtained by incorporating both the nuclei contours as seeds in a grey-scale 3D watershed approach based on raw images of actin. Lastly, the complete organoid contour is obtained directly from raw channels using fine-tuned thresholding and morphological mathematics filtering (all details in **Supplementary Figure 1** and **online methods**). Notably, these 3 different 3D segmentations are achieved within a remarkable timeframe of less than 2 minutes per organoid, without the need for GPU-based processing (using e.g. Intel(R) Core (TM) i7-8650U CPU @ 1.90GHz 2.11 GHz, 16Gb RAM). Segmentation processes of cellular structures have been validated in different acquisition mode (**Supplementary Figure2**) with a range of optical resolution and signal-to noise ratio, but all with the respect of the time limitation of one minute per organoid) described previously. As examples, we have shown that spinning disk acquisition (20X and 40X air objectives) as well as a point laser confocal (30X silicone objective) are compatible with our nuclei segmentation without any needs to retrain AI network by giving less than 10% of errors. This has been validated for a large range of signal-to-noise ratio (i.e. from 2.4 to 7.1) and of raw voxel sizes (**Supplementary Figure2a**). We also performed a quality control of our cell segmentation showing less than 8% of mis-segmented cells (**Supplementary Figure2b-c**). Eventually, this process generates an extensive list of hundreds of morphological and topological descriptors organized into an interpretable database establishing a robust foundation for detailed analysis.

We complete this pipeline by allowing biological interpretation of the results using a dedicated data mining work-flow, akin to cytometry analysis (**Supplementary Figure 3**). This interface enables the comprehension of the resulting database, the 3D segmentation layers, and the raw images. It empowers researchers to extract valuable biological insights and provide feedback on the raw images, facilitating the interpretation and utilization of the acquired data (**Supplementary Figure 4**). Some examples of data handling and navigation are provided in **Supplementary Video 1**. For clarity, we have called our data mining interface OrganoProfiler, but our database organization (described in **Supplementary Figure 3**) is fully compatible with already developed data mining open-source interface such as VTEA (https://imagej.net/plugins/vtea). Alternatively, users can import and generate similar graphical interface and statistical analysis using KNIME (https://www.knime.com/).

### 2/ Navigating through the 3 levels of segmentation, basic manipulations, image feedback and statistical analysis

These three levels of segmentation, graphical filtering, and image feedback provide a powerful framework for interactive exploration and analysis of complex biological data. Researchers can navigate through the data, manipulate visual representations, and apply filters to refine their analysis. We are providing examples of basic manipulations, image feedback and extractions of results in **Figure 2** based on cancer cell spheroids labelled with Phalloidin and DAPI. The graphical filtering or ‘gating’ provides researchers with the flexibility to exclude unwanted objects or artifacts from their analysis (**Figure 2a-b**) and the real-time feedback on segmented and raw images empowers users to make decisions during the filtering process, enhancing the accuracy and reliability of the results. The scatter plots revealed multi-scale correlations, such as the inverted relationship between the number of nuclei per organoid and mean nuclei volume. In our analysis of organoids with varying sizes (**Figure 2b**), we found for example an inverse correlation between the number of nuclei and the mean nuclei volume indicating that nuclei of larger spheroids tend to have a higher chromatin compaction rate (smaller volume) as compared to smaller spheroids. The segmentation and analysis of nuclei centroid distance from the spheroid border allow for the identification of spatial patterns and correlations within the spheroids (**Figure 2c**). This information provides valuable insights into the distribution and arrangement of cellular nuclei, aiding in the understanding of cellular behaviours within the spheroids. As example, we have applied a simple gating technique based on the distance of nuclei to the border. It has revealed that the external nuclei had a significant small increase in volume (+50 µm^3^, p<0.0001) compared to the internal nuclei. This finding suggests a relative relaxation of the mechanical constraint on the external nuclei.

**Figure 2:**
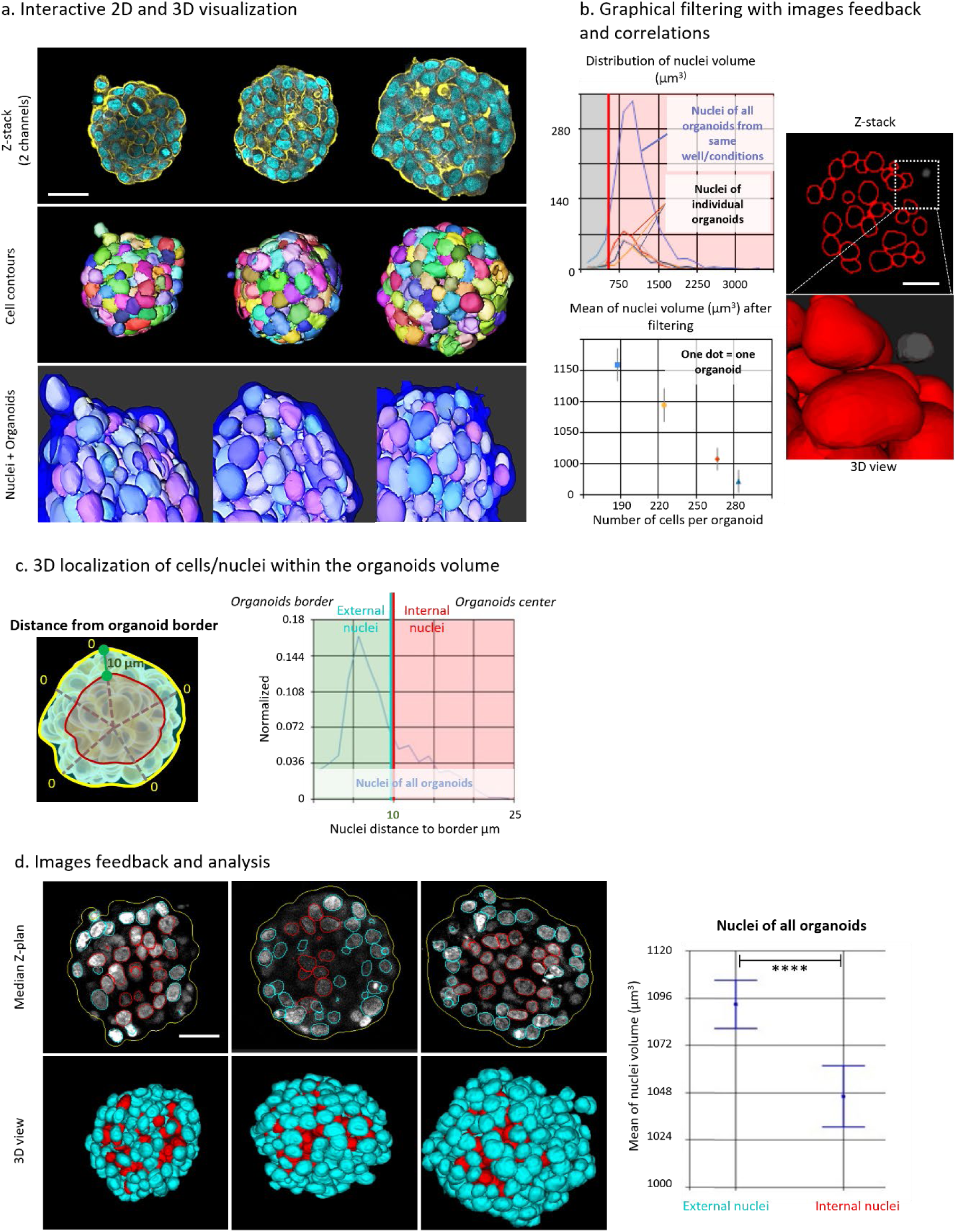
Interactivity and data navigation through three levels of 3D segmentation, graphical filtering, and image feedback. **a.** Examples of 2D and 3D visualization of cancer cell spheroids (HCT116) labelled with Phalloidin (gold) and DAPI (Cyan), showing 3D cell contours (multicolour, middle panel), and 3D nuclei contours (multicolour) with organoid contours (transparent blue, lower panel). Scale bar: 60 µm. **b**. Example of graphical display and filtering (gating) on a one-dimensional histoplot. The blue line represents nuclei volume of all cells in all organoids, while the other coloured lines represent nuclei of individual organoids. Manual filtering (red bar) can be adjusted to exclude smaller objects (e.g., debris or micro-nuclei), with feedback provided on the segmented or raw 2D and 3D images. The 2D scatterplot/or 2D-plot shows the number of cells per organoid for the four organoids versus the mean nuclei volume (after filtering, grey line for standard deviation). **c**. Scheme and histoplot of nuclei centroid distance from the external border of organoid. Filtering in two classes: green-cyan for nuclei within 0 to 20 µm and pink-red for nuclei at a distance greater than 20 µm. **d**. Image feedback in both 2D and 3D view. The boxplot displays the mean and standard deviation of nuclei volume (in µm3) for each class across all organoids. ****: p<0.0001, scale bar: 30 µm.

### 3/ Validation of Morphological Descriptors through Assessment of Well-Known Conditions of Morphological Modifications

We proceeded to validate the accuracy of our 3D segmentation and morphological descriptors by inducing morphological changes and compaction in cells using an osmotic stress (**Figure 3**). Osmotic stress, achieved here using a hypertonic medium, has been employed in previous studies as a means to simulate mechanical loading, resulting for example from a stiffer extracellular matrix (ECM) in the local environment^20^. Here, we used it as a positive perturbator of cellular morphology on cancer spheroids. Our findings have demonstrated that under hypertonic conditions (high-salt concentration medium), cells within spheroids exhibited a rounder morphology (**Figure 3a**). This was evidenced by a significant increase in the roundness of all cells highlighting by a clear alteration of cell roundness distribution. By applying plot gating, we observed under hypertonic condition a higher percentage (+24% as compared to isotonic condition) of cells within the ’High roundness’ gate. Notably, thanks to the direct image feedback capability of our method, we observed that the increased roundness is present for cells within the internal regions of the spheroids as well, thus indicating that the effect of the osmotic stress propagates through a dense 3D cell culture such as a cancer spheroid.

**Figure 3:**
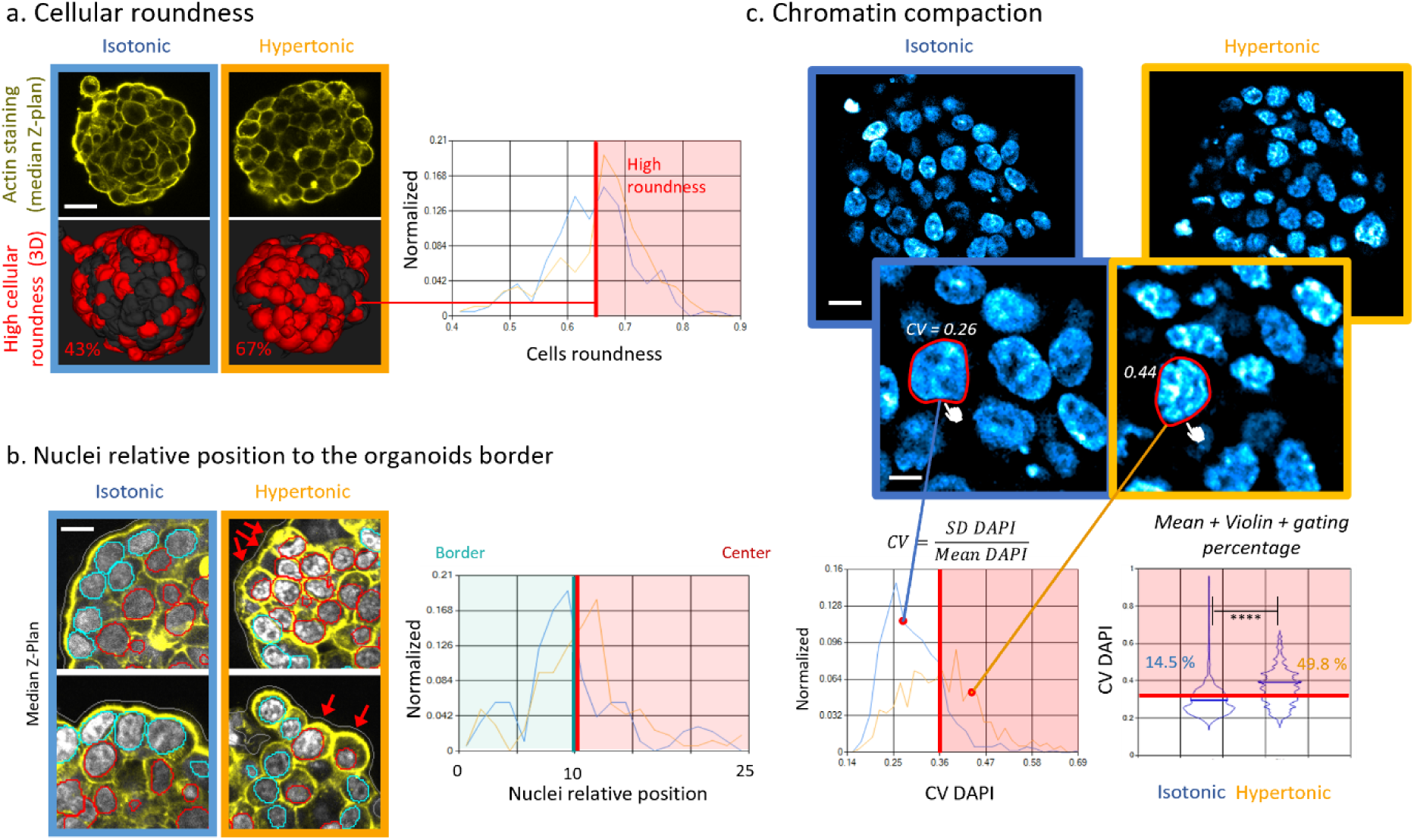
Proof of concept experiment on a well-known conditions of morphological modifications to validate multi-scale morphological parameters. **a.** The roundness of cells was compared under isotonic (blue) and hypertonic (orange) conditions. Representative images of cell shape from raw actin images in the median Z-Plan. Filtering was applied using a normalized histoplot with a red bar indicating the threshold of high cell roundness. The percentages of cells exhibiting high roundness are displayed on 3D segmentation (indicated by red). 2 organoids (223 and 284 cells) scale bar: 30 µm. **b**. Relative position of nuclei to the border of the organoid was compared under isotonic (blue) and hypertonic (orange) conditions. Representative images (2 distinct organoids per condition) of nuclei contour with cyan contours representing nuclei close to the border (within 10 µm) and red contours representing nuclei located farther from the border (>10 µm). Raw DAPI and Phalloidin images are shown in grey and yellow, respectively, in the median Z-Plan. Filtering was performed to classify nuclei into two classes (cyan and red) based on the threshold of 10 µm, applied to the normalized distribution of nuclei relative position to the organoid border. The red arrows indicate external nuclei belonging to the class of >10 µm, scale bar: 10 µm. **c**. Chromatin compaction was compared based on the calculus of the coefficient of variation (CV) of DAPI staining under isotonic (blue) and hypertonic (orange) conditions. Representative images of DAPI staining distribution (raw images) and zoomed images with examples of CV values displayed. The images/plots direct link is exemplified by red dots indicated the corresponding clickable nuclei contour (white hands). Filtering was applied (red bar) to identify nuclei with high CV values. Violin plots of CV values of the 2 conditions with the nuclei percentage in the ‘High CV’ gate (mean values represented by blue bars). ****: p<0.0001, analysis made on 10 organoids (5 organoids per condition,1080 and 1154 cells), scale bar: 15 µm and 5 µm for the zoom pictures.

Next, we looked for modifications of the 3D position of the nuclei relative to organoid’s border in the same hypertonic conditions (**Figure 3b**). As highlighted, a notable shift in the distribution of all nuclei positions towards the centre of the organoids can be observed (representative images) and quantified (significant increase in the median position of the external nuclei). These findings suggest that the applied osmotic shock induced changes in the spatial organization of nuclei within the organoids. This streamlines the analysis process by providing immediate visual representations of identified classes and gates.

In addition, our pipeline offers users the capability to identify effects through visual inspection of raw images and compile additional descriptors to confirm or refute initial visual impressions. For instance, by visual inspection, we observed first a change in the granularity of the DAPI staining under osmotic pressure. Nuclei from all organoids exposed to the hypertonic medium exhibited a distinctly more punctiform DAPI staining compared to the control conditions (**Figure 3c**). This change is commonly described as a chromatin compaction effect and can be quantified by calculating the coefficient of variation (CV) of the DAPI staining^21^. Our interface enables users to perform calculations across all features in the database (**Supplementary Figure 4**), allowing for the generation of new parameters to quantitatively confirm visual impressions. Users have the flexibility to enrich the database with additional features of their choice. In this case, we validated our visual inspection findings by quantifying the modification of the distribution of DAPI staining with the CV. The control condition displayed a normalized distribution with a narrower gaussian shape, whereas the hypertonic conditions exhibited a dispersed normalized distribution. Furthermore, we observed a significant increase in the mean CV (+30%, p<0.0001) and a higher percentage of nuclei into the ‘high CV’ class (from 14.5% in control condition to 49.8% in hypertonic medium).

By inducing morphological changes, we were able to validate the accuracy of our 3D segmentation and morphological descriptors. The observed increase in cell roundness, accompanied by changes in the nuclei positioning and staining granularity, provides robust evidence of the effectiveness of our approach in capturing and quantifying cytoplasmic and nuclear morphological alterations.

### 4/ Cell-to-Neighbourhood 3D topology descriptors to identify tissue patterning and effects of extracellular microniche

Within organs and tissues, cells intricately interact to create layers, tubes, or complex clusters that possess a distinctive three-dimensional organization (tissue patterning), which is crucial for the proper functioning of the organs. Furthermore, it is widely recognized that local topological features (e.g. curvature) of the extracellular microniche plays a pivotal role in facilitating the appropriate differentiation of cells by modifying morphology and cell-cell relative position^8^. Analysing the tissue patterning and thus cell relative positions is essential for understanding these differentiation and homeostasis processes in 3D. In our previous work, we successfully applied an AI strategy using a dedicated pretrained U-net network (as referenced in Beghin et al.^22^) to identify and three-dimensionally segment neural rosettes, complex internal arrangements specific to neuroectoderm organoids. However, it is important to note that this approach has limitations. First, a substantial number of organoids, often over a hundred, are required to build the 3D imaging database used for training. Moreover, creating manual ground truth contours to train the network for this particular structures not only consumes a significant amount of time but also necessitates the involvement of multiple experts to achieve accuracy and consensus. Eventually, the specialized U-net network we employed is inherently tailored exclusively to neural rosettes structures, characterized by specific staining and image quality. It lacks versatility and necessitates prior knowledge of the expected target structure. To address all these limitations, we propose an innovative and versatile solution based on the Cell-to-Neighbourhood relative position, using only the 3D nuclei positions without any needs of other markers. 3D topology descriptors were compiled based on the 3D centroid positions of nuclei, as described in **Supplementary Figure 5** and the **Online Methods** section. In brief, these descriptors not only quantify the intricate cell-to-cell positions (distance and density) in the 3D cell culture but also provide precise measurements of neighbouring nuclei distribution by fitting ellipse around the surrounding nuclei of each nucleus and quantify this ellipsoidal shape by axis measurements and their ratio (See **Online Methods** and **Supplementary Figure 5**). We conducted experiments using normal primary epithelial breast cells (HME cells) seeded in a cup-shaped microniche composed of extracellular matrix (ECM) as proof-of-concept experiment.

Here, the specific cup-shaped microniche structure forces cells to position themselves in a typical, non-random distribution, forming an adherent monolayer on different types of planes (from the top to bottom: long vertical curved walls, then short diagonal curved planes and, at the bottom of the ECM cup, a restricted horizontal plane) (**Figure 4**). To measure this multicellular organization in 3D, the ellipsoid fitting was performed on the neighbouring nuclei of each nucleus, with the ellipsoid’s three axes obtained for each nucleus (**Figure 4a**). Using the ellipsoid measurements, ‘Prolate’ and ‘Oblate’ ratios were calculated, which were then plotted in a 2D scatterplot to reveal patterns of 3D organization and enable multi-gating processes.

**Figure 4:**
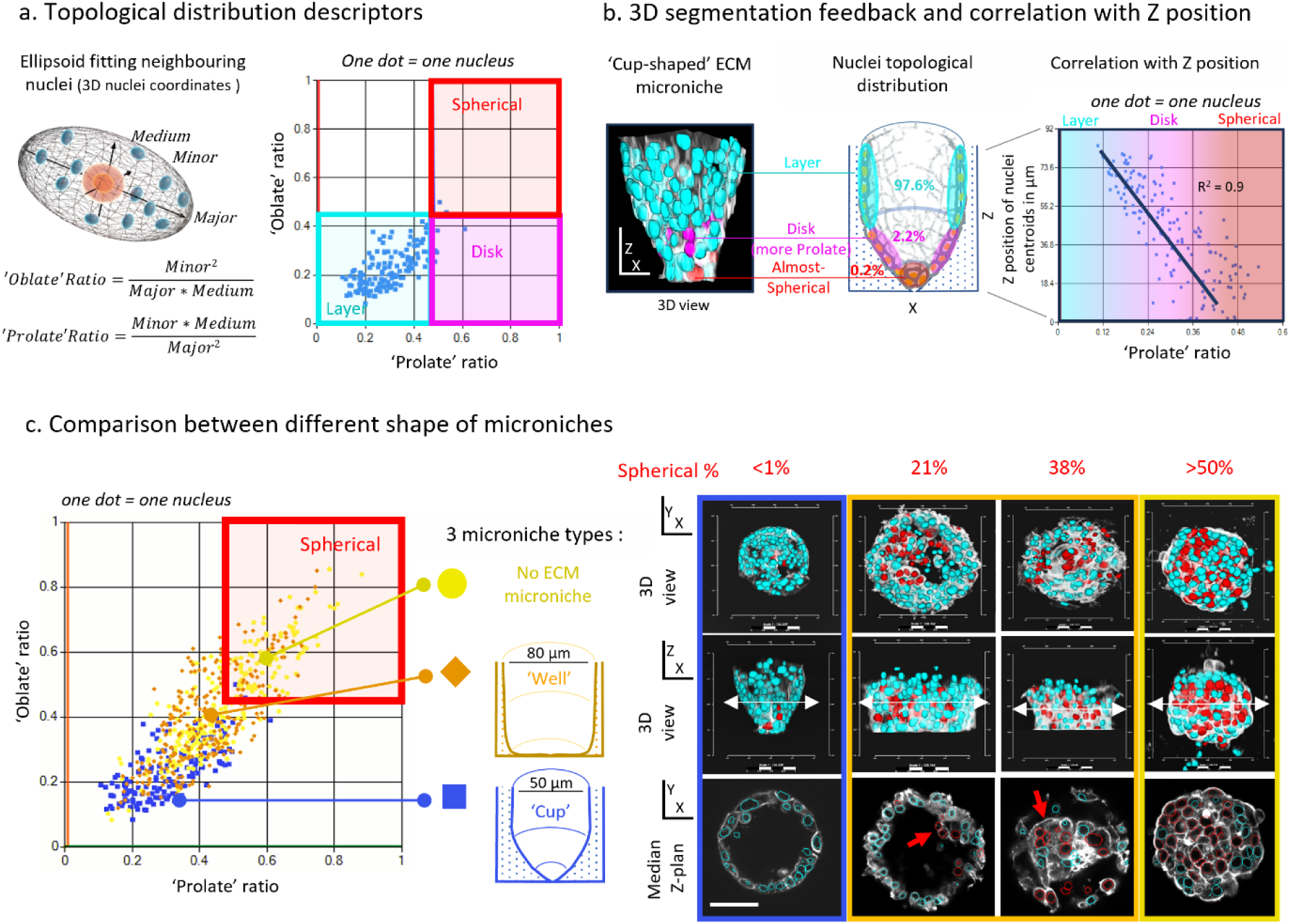
3D topology descriptors and effects of microniche shaping on cellular 3D positioning within microphysiological system. **a**. Ellipsoid fitting of neighboured nuclei scheme with axis of the ellipsoid and calculation of Oblate and Prolate ratio. Scatterplot of Oblate ratio vs. Prolate ratio with multi-class gating of a ‘cup-shaped’ ECM microniche seeded with of human mammary epithelium cells (each dot representing one nucleus). **b**. Color-coded feedback of the 3 classes on the corresponding 3D segmentation, percentage of nuclei of each class represented on a schematic view of the cup-shaped microniche. Scatterplots and correlation with Zposition with either Prolate or Oblate ratio (blue line: linear regression, R^2^). **c.** Topological comparison between cells seeded on 2 different shapes of microniches (Blue: ‘Cup’, Orange: ‘Well’) and no ECM microniches (Yellow) on a scatterplot Oblate vs. Prolate ratio (one dot = one nucleus). Color-coded feedback (red contours for ‘Spherical’ gate, cyan for all other nuclei) on 3D segmentation views (XY and XZ) and on a median XY plan (corresponding to the white double-arrows) with percentage of nuclei with ‘Spherical’ ellipsoid for each microniche (corresponding to an isotropic neighbourhood). Red arrows highlight anarchic multilayers of cells. Scale bar= 25 µm.

In a first example, three gates were defined based on the ellipsoid characteristics: the "Spherical" gate represents nearly spherical or isotropic shapes (and in our microniche, 0.2% of nuclei population), the "Disk" gate represents flattened shapes (2.2% of nuclei), and the "Layer" gate corresponds to elongated and flattened shapes (97.6% of nuclei). Applying these gates with color-coded feedback on the 3D segmentation, we observed the expected correlation between the ellipsoid types and the microniche planes along the principal axis of the microniche (Z-axis) (**Figure 4b**). Correlation analysis through 2D scatterplots of Prolate ratio versus Z-position of nuclei revealed a linear regression (with R^2^=0.9) between the Prolate ratio and Z-position (**Figure 4b**). By applying these analytical approaches, we successfully validated the correlation between our Cell-to-Neighbourhood 3D topology descriptors and microniche surface, providing insights into the 3D organization of cells within the microniche.

Then, we used the same approach to compare two types of microniche configurations: the previous ECM cup-shape with a 50 µm diameter (surfaces completely cell adhesive) and a flat-bottom well-shape with an 80 µm diameter displaying only 1 type of adhesive plane: long vertical curved walls and a large flat horizontal non adhesive surface, along with a spheroid without ECM, meaning non adherent surface available for the cell (**Figure 4c**). For clarity, we focused on presenting the percentage of “Spherical” ellipsoid configuration. The results demonstrated clear differentiation among the microniche types, and the spheroid configuration based on the percentage of “Spherical” ellipsoid (indicative of isotropic/random cell-cell positioning). The 50 µm cup-shaped microniche exhibited a lower percentage (<1%) of randomly positioned cells compared to the 80 µm flat-bottom well, which showed a range of 20 to 40% of nuclei randomly positioned with some anarchic multilayer patterns (**Figure 4c**). In contrast, a typical cancer spheroid without ECM microniche presented a percentage of randomly positioned cells higher than 50%. These findings, derived from advanced topology descriptors, confirm that the precise shaping and size of the ECM microniche strongly influence the 3D organization of cellular patterns. Other descriptors such as local densities and mean distances (Cell-to-Cell) between nuclei are also provided (**Supplementary Figure 5b**).

The analysis described above provides valuable insights into the Cell-to-Cell and Cell-to-Neighbourhood spatial relationships and sheds light on the impact of microniche shaping the multicellular organization. This approach is highly pertinent for examining the internal organization of 3D cell culture structure, specifically to determine if they exhibit non-random cellular arrangements mirroring native tissues, all without the need for prior knowledge, additional segmentation steps, or AI training.

### 5 Unsupervised/supervised workflow to automatically extract signatures from a ‘discovery’ experiment

Within this section, we introduce an elevated echelon of automated analysis, thereby integrating methodology that enlists both unsupervised and supervised modalities. This methodology is dedicated to the extraction of distinct hallmark attributes (=signature) concomitant with three-dimensional (3D) morphological characteristics. Its implementation is demonstrated in an experiment characterized by its exploratory nature, conducted during a parabolic flight (**Online Methods**). Briefly, hundreds of breast cancer spheroids were placed aboard on an aircraft performing a succession of 15 parabolas and fixed whether onboard after the last run of the 15^th^ parabola (Day0, F for Flight) or after 24 hours (Day+1, F24 for Flight fixed after 24 hours). Corresponding Ground Controls have been cultivated in the laboratory and fixed at the corresponding same time (GC and GC24). This set of extreme conditions simulates a regimen of cyclic mechanical stresses involving fifteen parabolic trajectories, each demarcated by distinct phases of accelerations of 2.5g, transitions to simulated microgravity (0g), and reverting to 2.5g forces.

It’s worth highlighting the significant absence of prior results obtained in such comparable conditions. This void is particularly striking given the contemporary surge of interest in space biology and the concurrent application of organoids in space-related extreme conditions^23–25^. Earlier deployments of biological experiments in parabolic flight have yielded some first insights into the immediate effects on cellular adhesion but only within a 2D cellular monolayer^26–28^. The dataset used here is thus firmly situated within the methodological framework of inquiry driven by discovery. This approach is underpinned by the overt lack of prior knowledge pertaining to the specific morphological traits of cells within a 3D tissue-like structure, that might undergo alterations under the influence of the mentioned cyclic mechanical stresses.

The contingency involved 120 stacks of organoids, which were subjected to the described conditions (**Figure 5a**) (Ground Control (GC) and Flight (F) on Day 0, Ground Control 24 (GC24) and Flight 24 (F24) on Day+1). These spheroids were digitized using our streamlined method, resulting in an extensive collection of 3D morphological features from more than eight thousand cells. Importantly, the cells were numerically pooled ‘blindly’ or anonymously, setting aside the identifiers related to the conditions and to organoids. This ‘blind’ pool was then utilized for unsupervised techniques, such as Principal Component Analysis (PCA). This technique was employed to effectively reduce the dimensionality of the data to its primary component, PC1. Subsequently, cells with all their identifiers based on their respective conditions were plotted according to their PC1 values. This visualization strategy brought to light distinct signatures within the data and specific cell populations can be isolated using gating on PC1 value. Interestingly, immediate visualization from this clustering was made available through our comprehensive interface, allowing for feedback visualization in both 2D and 3D image formats (**Figure 5a**). The visualization indicated a noticeable concentration of the ’High PC1’ population along the outer periphery of each organoid. This immediate 3D localization of a particular cell population based on a multiparametric signature serves as a guidance to biological meaning. For example, here, users can apply manual gating strategies, like using nucleus distance from the border, to isolate external cell population and then inspect individually the morphological features. In the case of the data reduction analysis (e.g. PCA but not restricted to), the user does not have any a priori knowledge about a specific localization of the cells of interest nor the type of effect expected. Initially, the user blindly generates a cell signature, and our interface allows raw images feedback and 3D localization, thus biological interpretation. In a classical data mining approach, the user first applies filters and strategies based on his/her expertise and prior knowledge of the suspected effects. Subsequently, the user generates statistics that either support or refute his/her initial hypothesis. But this user-based approach can be very high time consuming, quite limited or can even never reveal any significant differences between conditions in a case of such discovery like experiment.

**Figure 5:**
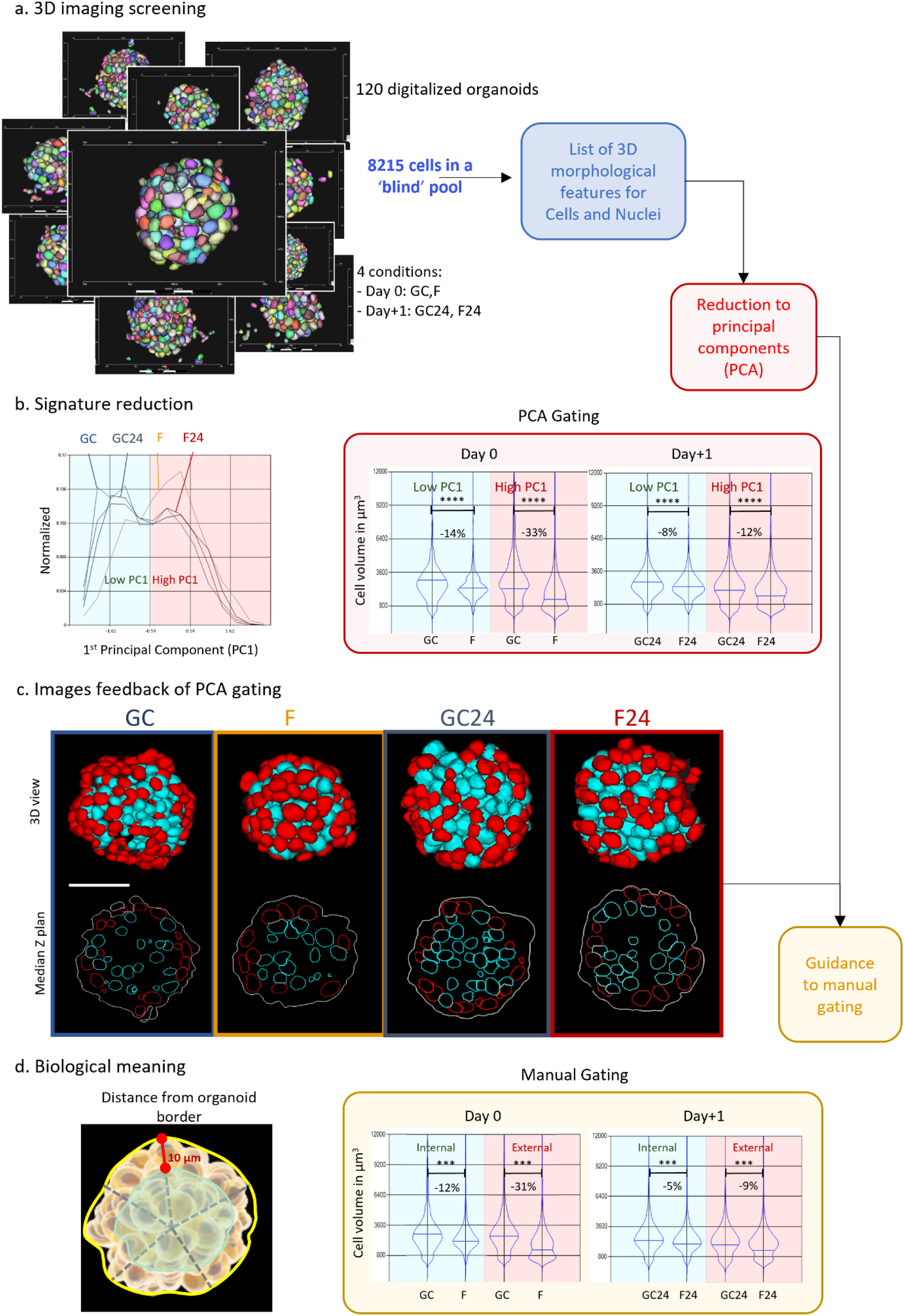
Integrated Workflow for 3D Morphological signature understanding. **a.** An automated 3D imaging screening generated 120 stacks of organoids under four conditions (Day 0: Ground Control (GC) and Flight (F), Day+1: Ground Control (GC24) and Flight (F24)). These organoids were fully digitized, resulting in numerous 3D morphological features across over eight thousand cells collectively amalgamated without condition descriptors (‘blind’ pool). **b.** This ‘blind’ pool is employed for unsupervised techniques, notably Principal Component Analysis (PCA), to reduce the data’s dimensionality to the first principal component (PC1). The cells, now identified by their respective conditions, were then plotted based on their PC1 values. This visualization highlighted distinct signatures, and subsequent clustering isolated specific cell populations (Red: high PC1 value, Green: low PC1 value). Violin plots represent unsupervised clustering of PC1, illustrating differences in cell volume among conditions (Day 0: GC and F, Day+1: GC24 and F24, blue bars indicate means, ****: P<0.0001). **c.** Immediate images feedback of this clustering, available in both 2D and 3D view through our comprehensive workflow, showcased a prevalent localization of the ’High PC1’ population (red) at the outer border of each organoid. **d.** Subsequent biological meaningful strategies based on images feedback, such as applying filters based on nucleus distance to the organoid’s border. Yellow plots of guided filtering on the nuclei distance to the border with significant differences of cell volume (blue bars for mean, ****: P<0.0001).

Here the two strategies revealed that cells have undergone a decrease of their volume due to the exposure to the parabolic flight (**Figure 5b-c**). This significant strong diminution was maximized for cells of the High PC1 population as compared to the Low PC1 population at Day 0 (-33% and -14% respectively). Less but significant decreasing of cell volume was also obtained at Day+1 (High PC1: -12% and Low PC1: -8% respectively) (**Figure 5b**). The user-based approach was able to generate same profiles of response with similar significant decreasing of cells volume, and same difference of response correlated with cells position to the external border (**Figure 5C**). We can here conclude that the morphological stress generated by parabolic flight on spheroids affect more importantly the morphology of external cells (=border of the spheroids) and that this differential effect is still significantly observed after 24hours in normal G condition of culture. Furthermore, this effect is impacting not only on cell volume but also on other features such as cell density and nuclei volume (**Supplementary Figure 6**).

## Discussion

Overall, our approach demonstrates a powerful and efficient methodology for imaging, analysing, and extracting biological information from 3D cellular microsystems. By combining and adapting state-of-the art machine learning with classical image analysis and common 3D microscopy techniques within the respect of biological constraints, we here aim to reveal precise morphological modifications of nuclei, cells and of 3D culture itself. We have developed quantitative metrics to better understand the complex interactions that occur at the cellular and multicellular level. As highlighted, it encompasses various representations, plots, filtering, and summary procedures, streamlining the analysis process and enhancing user efficiency. Moreover, by employing advanced topology descriptors like the Cell-to-Neighbourhood and 3D segmentation feedback, we can evaluate and quantitatively assess the degree of organization and similarity between organoids and their corresponding tissues. Our method is additionally founded on the integration of widely employed methodologies but from entirely distinct biological technologies. These methodologies encompass cytometry-like gating and Next Generation Sequencing data reduction approaches with the originality to allow feedback on 3D or 2D raw images and segmentation meshes. Our innovation introduces a groundbreaking method for generating comprehensive 3D cell morphological signatures within organoids, offering a transformative way to assess different mechanical constraints.

The set of proof-of-concept experiments displayed here shows the reliability of our methodology in characterizing 3D morphological changes in complex multicellular structures such as organoids or spheroids. A simple set of few organoids or large batches of thousands of organoids in a multiwell plate can be analysed using our method. Furthermore, as whole organoid segmentation is not mandatory for generating the database of cell and nuclei descriptors, our method can also be applied to analyse other cell culture systems, such as monolayers of cells on flat or curved surfaces. However, our segmentation method does require nuclei segmentation, necessitating a DNA staining procedure independently of any specific fluorescent wavelength used. Cell segmentation can be accomplished using either actin staining or, if necessary, a dedicated membrane staining. In the context of phenotype analysis, which involves evaluating the expression levels of proteins, measuring other specified fluorescent intensities is a straightforward process with intensity-based values such as mean, minimum, maximum, median, and standard deviation.

This analytical framework enables a comprehensive investigation of the structural and cellular characteristics of 3D cell culture system or tissues, enhancing our understanding of their fidelity and potential applications in tissue engineering and injury. By transcending differentiation-driven variations, our approach allows for a comprehensive understanding of how cells respond to external cues and perturbation. The quantification of 3D cellular deformations within intricately designed tissue-mimicking structures represents a significant contribution to the advancement of our comprehension of cellular biomechanics, tissue homeostasis, and therapeutic interventions within the biomedical domain. Moreover, this innovative approach holds substantial promise for applications in more extreme conditions, such as those encountered in the field of space biology. By enabling a comprehensive understanding of how cells respond to mechanical constraints within both controlled laboratory environments and extreme settings, this methodology opens new avenues for research and the development of novel compounds.

## DATA AND MATERIAL AVAILABILITY

Complete dataset including 3D raw images, segmented images and database will be available upon request. We will provide access to our interface for testing based on Material Transfer Agreement.

## ACKNOWLEDGMENTS

AB acknowledges the funding support of MBI seed grant and the Singapore EDB’s Office for Space Technology and Industry (OSTIn). AB and GG thank C. Fischer (campaign manager) and the members of Prof. Ullrich team for providing organizational and logistical support (University of Zurich). All authors acknowledge the Royal Netherlands Aerospace Centre (NLR) and TU Delft, especially Arun Karwal, Theodoor Johan Mulder, Hendricus De Haan, Gijsbert Frederik De Toom and Marcel Johan Verbeek for the flying parabolic flight campaign. We thank the Swiss SkyLab Foundation for the operative organization of the 6th Swiss Parabolic Flight Campaign. AB and GG thanks H.Y. Bong for sample preparation and shipment for the parabolic flight. All authors thank A. Wong and D. Pitta de Araujo for their help editing the manuscript.

## ONLINE METHODS

### Maintenance of Cell culture and medium

Colorectal cancer cells HCT116 (91091005, Sigma-Aldrich) and breast cancer cells MCF7 (86012803, Sigma-Aldrich) were cultured in DMEM high glucose (11965092, Gibco) supplemented with 10% Foetal Bovine Serum (FBS) (10082147, Invitrogen), 100Uml−1 penicillin-streptomycin (15070063, Invitrogen) at 37 °C and 5% CO2 (=complete medium).

HME cells (Human Mammary Epithelial cells, purchased from ATCC, PCS-600-010), primary mammary epithelial cells, were cultured in complete Mammary Epithelial Cell Basal Medium (PCS-600-030, ATCC) supplemented with Mammary Epithelial Cell Growth Kit (PCS-600-040, ATCC). HME cells were subjected to no more than ten passages in culture when used in experiments.

### Formation of spheroids

After trypsinization, the cell lines (HCT116 cells or MCF7 cells) were suspended in complete medium and adjusted with a concentration of 0.3 × 10^6^ cells/mL. Next, 1ml of cell suspension was dispensed in p24 well (5826-024, Iwaki) plates containing JeWells that were already undergone long-term passivation with 0.5% Lipidure (CM5206, NOF America), as previously described^22,29^. The cell culture plates were placed into the cell incubator (37 °C, 5% CO2, and 100% humidity) for 5 min to get approximately 20-50 cells per spheroid. The cell suspension was removed, washed three times with DPBS (14190250, Invitrogen) and 1ml of complete medium was added per wells. The medium was changed every 3 days. The cells were fixed after 1 week of culture for staining.

### Osmotic stress treatment

After 1 week of HCT116 spheroids formation as described above, osmotic stress was applied by adding NaCl into the culture medium as previously described ^30,31^. Complete medium supplemented with 100 mM NaCl (7647-14-5, Merck) was used as hypertonic medium. Complete medium of HCT116 spheroids was removed and replaced with the hyperosmotic medium for a 6 hours incubation in 5% CO2 at 37 °C, followed by fixation for staining.

### Microniche-shaped extracellular matrix

Microniche-shaped extracellular matrix (ECM) arrays were created using elastomeric stamps [PDMS, poly(dimethylsiloxane)] containing the desired cup-shaped (50 µm diameter) or flat-bottom (80 µm diameter) pillars. Briefly, the PDMS working mold was produced as a replica cast of a primary mold, following standard soft-lithographic procedures. We used Sylgard 184 (Dow Cornig), prepared by thoroughly mixing and outgassing the base resin an its reticulation agent in 10:1 weight ratio. Then the mixture was poured on the primary mold and degassing in a vacuum jar was used a second time to ensure complete filling of the cavities and removal of any trapped air. The PDMS was then thermally cured for 1-2 hr at 65 °C on a Hot Plate, finally after cooling down the PDMS was peeled off and diced to final dimensions for usage.

In 35mm glass bottom dish, the PDMS working mold with pillars of defined size and shape was pressed to a hydrogel layer of Methacrylated Gelatin (GelMA) containing photoinitiator lithium phenyl-2,4,6-trimethylbenzoylphosphinate (LAP) (5272, Advanced Biomatrix) at 50 °C. Next, GelMA-LAP was crosslinked using UV-LED box (UV LED KUB2, Kloe France, 365nm) with a power density of 35 mW/cm2, for 1 min and the mold was carefully peeled off. Microniches were then covered with DPBS and place in the cell incubator.

### Primary cells seeding on ECM microniches

After trypsinization, HME cells were suspended in complete Mammary Epithelial Cell Basal Medium containing 5 µg/ml rh-insulin, 6 mM L-Glutamine, 1 µg/ml Epinephrine, 5 µg/ml Apo-Transferrin, 5 ng/ml r-H-TGF-α, 0.4% ExtractP, 100 ng/ml Hydrocortisone Hemmisuccinate and adjusted with a concentration of 0.3 × 10^6^ cells/mL. Then, 2ml HME cells were seeded at the top of microniches GelMA-LAP crosslinked. The cell culture dish was placed into the cell incubator (37 °C, 5% CO2, and 100% humidity) for 5 min to allow cells to enter microniches cavities allowing approximately 20-50 cells per microniche. The top cell suspension was removed, washed three times with DPBS (14190250, Invitrogen) and 2ml of complete Mammary Epithelial Cell Basal Medium was added into the 35mm glass bottom dish. The medium was changed every 2 days and the cells were allowed to self-organize into curved layers or epithelial cell adopting the geometry of the hydrogel microwells. The cellular microniches were fixed after 2 weeks of culture for staining.

### Parabolic flight

The parabolic flight experiments were carried out on-board of a Cessna Citation II (Royal Netherlands Aerospace Centre (NLR)), organized by the Swiss SkyLab Foundation and the UZH Space Hub (Innovation Cluster Space and Aviation of the University of Zurich) as part of the Swiss Parabolic Flight Program^32^. During a parabolic flight manoeuvre, 15 consecutive parabolas occurred in three steps. Firstly, in the pull-up phase, the nose of the aircraft is raised to about 60 degrees for 20 seconds (hypergravity at 2.5g). Then, in the second phase, the aircraft is in free fall and experiences microgravity for 13 to 15 seconds following a parabola trajectory. Finally, in the pull-out phase, the aircraft is tilted down at approximately 60 degrees for another 13. 15 seconds (hypergravity at 2.5g). During a parabolic maneuver, an aircraft is weightless due to flying on a Keplerian trajectory, an unpropelled body in an ideally frictionless space that is subjected to a centrally symmetric gravitational field^33^.

### Cell culture procedure for the Parabolic flight

Before the parabolic flight, MCF7 spheroids were cultured, in a tissue culture room close to the airfield, in p24 well plates (n=4) containing JeWells for 1 week as described above. Two p24 well plates (n = 2) were used as 1 g ground control. The other two well plates were taken on board of the aircraft for the parabolic flight using a portable cell incubator (allowing normal cell culture conditions during all the transportation and flight: 5% Co_2_, 37°C, Cellbox). Ground control plates were fixed at the same corresponding time as the plate on-board and using exactly same procedure as described in the fixation procedure below. We referred to ‘GC’ and ‘F’ for the Ground Control and Flying plates that were fixed just after the last 15^th^ parabola (at day 0). ‘GC24’ and ‘F24’ corresponds to same conditions but fixed 24 hours after the flight (at day +1). All plates have been sealed using CO2 permeable polymeric films to avoid any medium lost during transportation and flights. After fixation, the plates were washed 3 times with DPBS and stored at 4 °C for staining and image acquisition.

### Fixation and staining for imaging

Spheroids and microniche cell layers were fixed for 15min in 4% paraformaldehyde (28906, ThermoFisher Scientific) at room temperature. Then the 3D cultures were permeabilized for 1 hour in 0.2% Triton-X-100 (T9284, Sigma-Aldrich) solution in sterile DPBS (14190250, Invitrogen) at room temperature on an orbital shaker. Samples were then incubated with 0.5μgml−1 DAPI (62248, ThermoFisher Scientific) and Alexa Fluor 647 phalloidin (A12379, ThermoFisher Scientific, 1:200) at room temperature for 30min on an orbital shaker followed by three rinsing steps with washing buffer before image acquisition.

### Image acquisition process

The images of spheroids were obtained with a confocal laser scanning microscope (FV3000, Olympus, Tokyo, Japan). Excluding images acquired on a spinning disk confocal used for the quality assessment of nuclei (see below), all images were acquired using a 30x/1.05, WD 0.8, Silicone objective, with a z-step set to 3 μm, and XY resolution of 0.3*0.3 µm. Confocal images of DAPI (laser excitation 405 nm) and Phalloidin (laser excitation 640 nm) were acquired in less than a minute per spheroid. Multi-Area-Time-Lapse (MATL) automatic acquisition mode was used to image hundreds of spheroids per well.

### Image preprocessing using histogram matching method (optional)

The acquired 3D images were subject to intensity decay along z direction, necessitating intensity enhancement, particularly in the later z slices. To perform histogram matching using Skimage^34^, we calculated the 99th percentile intensity for each slice and selected the slice with the highest 99th percentile as the reference slice. Based on the reference slice, a histogram matching procedure was performed for all slices. This preprocessing was carried out for all channels.

### Nuclei segmentation by 3D Stardist

Nuclei segmentation was performed using 3D Stardist^15^ model CNN on DAPI channel. Our pipeline uses a model trained internally on realistic simulated 3D image data with a voxel size of (0.8μm, 0.8μm, 1μm).

The critical training phase was performed internally using realistic simulated 3D dataset. Generating synthetic data involves several sequential steps, starting with creating a labelled mask, where each voxel gets a unique integer identifier for specific nuclei. To achieve a diverse representation of nuclear morphologies, reflecting the complexity found in real-world scenarios, we integrated a combined strategy involving a parametric sphere representation coupled with a 3D deformation model^35^. The resulting labelled mask was considered as the synthetic ground truth. To produce the corresponding image, voxel values of this mask are transformed using techniques that mimic the acquisition procedures commonly employed in microscopic systems. We employed Perlin noise to replicate textural variations^36^ and Gaussian blur to emulate point spread function (PSF)^37^, introducing subtle imaging imperfections. In the end, through the manipulation of a range of simulation parameters, we have produced a dataset consisting of synthetic 3D images paired with labelled masks. This dataset has served as a valuable resource for training and evaluating various segmentation algorithm.

Notably, the generated dataset was constructed with a unique voxel size of 0.8µm* 0.8µm* 1µm. In order to fit the resolution of the trained network, users can use auto resize function for resizing their 3D data to match the voxel size of the training images. Alternatively, data can also be resized by manually input (xyz) resampling factor. To speed up the processing, we cropped our images to 512×512 pixels prior to segmentation. In our implementation, results of our 3D Stardist trained network can be optimized by fine-tuning several thresholds, such as the probability threshold (typically ranges from 0.3 to 0.7) to retain only entities with a probability exceeding this threshold, as well as the Non-Maximum Suppression (NMS) threshold (typically ranges from 0.1 to 0.3), which selects a single entity among overlapping entities. In our approach, size thresholds can also be set to filter out debris.

### Cell segmentation by seeded watershed

Cells were segmented by seeded watershed method of Skimage^34^. The nuclei segmentation results were used as the starting seeds. The fundamental principle of this algorithm involves identifying starting points, or “seeds”, typically associated with objects of interest in an image. These seeds serve as reference points from which segmentation propagates. The algorithm treats the image as a landscape and assigns pixels to the nearest seed’s "basin." In our context, the initial seeds for segmentation were derived from the nuclei segmentation results and their propagation was guided based on the intensity channel in which the cells were observed (i.e. Phalloidin). It has to be noticed that the cell expansion was confined within the organoid mask. Additionally, the expansion distance from the nucleus border to the cell border was limited to a maximum distance (which was set at 14μm by default but can be adjusted if needed). This prevents the over-expansion of cell from the nucleus border. Once the cell label mask was created, it was further refined using a 3D median filter with a sphere kernel.

### Organoid segmentation by Otsu method

Organoid segmentation was performed by Otsu method using mean intensity of all channels. Before segmentation, we enhanced contrast for all z slices by applying Contrast Limited Adaptive Histogram Equalization (CLAHE)^38^ preceded by image normalization. The contrast-enhanced images were then smoothed using a 3D Gaussian filter. Auto Otsu threshold was calculated using the stack histogram and the final threshold can be adjusted using a scale factor, which was set at 0.8, by default (i.e. *final_threshold* = *scale_factor* x *auto_otsu_threshold*). After thresholding, small elements were automatically filtered from the mask, to retain only the largest one in case of out-of-organoids debris or cells. To generate a smooth 3D mask, we conducted a series of mathematical morphological operations on a slice-by-slice basis, i.e. dilation with a disk structure element followed by hole filling and erosion with a disk structure element.

### Quality assessment of nuclei and cell segmentation

We conducted a thorough quality assessment of nuclei and cell segmentation using three distinct imaging configurations. The first configuration involved a Laser Scanning Confocal (LSC) FV3000 Olympus microscope equipped with a 30X silicone immersion objective featuring a numerical aperture (N.A) of 1.05 and a raw voxel size of XYZ 0.31*0.31*3µm. The second configuration utilized a Spinning Disk Confocal BC43 Andor system with a 20X air objective (N.A=0.8) and a corresponding raw voxel size of 0.325*0.325*1.5µm. The third configuration employed the same spinning disk system with a 40X air objective (N.A=0.75) and a raw voxel size of 0.1625*0. 1625*1µm. The Signal-to-Noise S/N ratio serves as an indicator of image clarity and the degree of noise interference, with higher values denoting superior quality (**Supplementary Figure 2a**). We calculated the percentage of missing or inadequately segmented nuclei, referred to as "% errors,". In each image, an expert counted the nuclei that were missing and those with a significant precision issue in their contours, which exceeded 30% of the total nucleus area. We did the same approach for cell segmentation between different Z-plane of the same organoid (**Supplementary Figure 2b**) and between different organoids at the same Z-plane **(Supplementary Figure 2c).**

### Extraction of morphological and topological descriptors

For each detection (nuclei, cell, organoid), a comprehensive list of morphological features such as volumes, elongation, roundness, equivalent ellipsoid diameters, bounding box sizes, centroid coordinates, distance to organoid border (for nuclei and cell) and intensity statistics for each channel (minimum, maximum, mean, standard deviation) has been generated. Other than these morphological features, a density analysis tool was also developed to extract 3D topological descriptors from nuclei position and organoid mask. The 3D topological descriptors are classified into nuclei spatial arrangement descriptors and nuclei density descriptors (**Supplementary Figure 5**). Cell-to-Neighbourhood (nuclei spatial arrangement) descriptors enable quantitative measurements of cell organization by fitting an ellipsoid from all nuclei coordinates in a spheroid of radius r (µm)(**Supplementary Figure 5a**) that can be specified by users. Ellipsoid’s axes length, prolate ratio, oblate ratio and various descriptors extracted from the fitted ellipsoid provide insights into internal organization of organoid such as tissue patterning. For Cell-to-Cell density analysis, nuclei density descriptors computed include distances between nuclei, local density, Ripley’s estimated number of nuclei in a spherical volume of user-defined radius, r (µm)(**Supplementary Figure 5b**).

### Data mining and unsupervised analysis

The complete workflow from segmentation (called OrganoSegmenter (OS) step) to data mining (called OrganoProfiler (OP) step) is shown in **Supplementary Figure 3**. Briefly, the 3D segmentation masks in TIFF files and database in CSV file, together with the multi-channel images (in composite or separate TIFF files), can be used for interactive data exploration and data mining (**Supplementary Figure 4**). Software dedicated for image-based data mining such as VTEA^39^, CellProfiler Analyst^40^ or KNIME^41^ provides important features such as gating, images feedback in 2D and 3D and statistical analysis. We have incorporated all these well-known functionalities with various others tools such as generating additional features to enable advanced data analysis. In our analysis, we created new feature, named as Coefficient of Variation (CV) by dividing standard deviation with mean intensity. The Principal Component Analysis (PCA), which is an unsupervised technique that can be used for dimensionality reduction for large database has been used to generate morphological signatures. In addition to highlighting the most meaningful features, this method is well known for allowing better visualisation which can aid in data exploration and understanding, noise reduction (as PCA focus on the most informative dimensions), generalization and computational efficiency for further classification^42^.

---

**Supplementary Figure 1:**
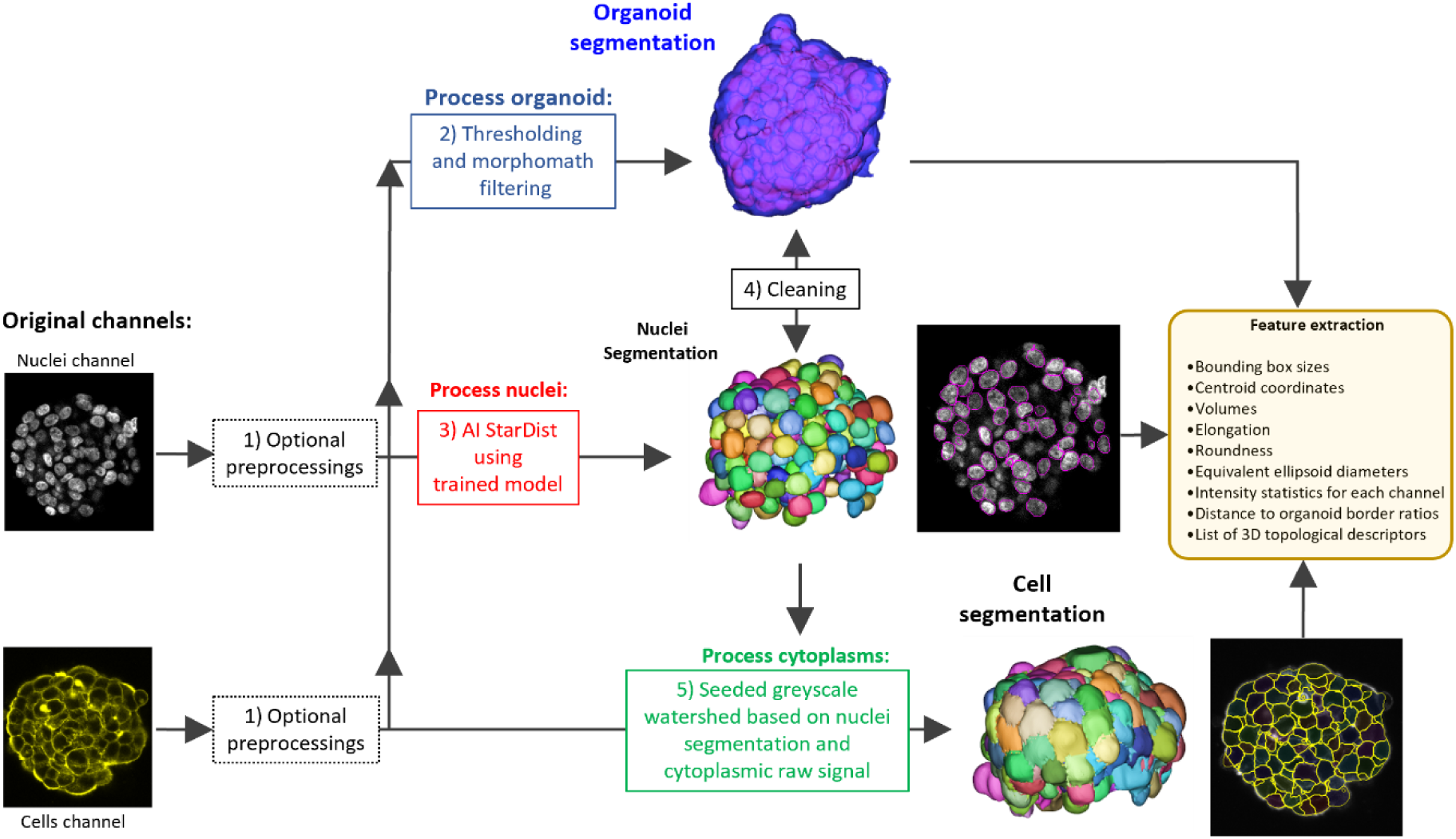
Scheme of preprocessing, segmentation processes and final results with features extracted. The pipeline starts from **(1)** optional preprocessing, e.g., histogram matching to correct intensity loss due to depth penetration. The preprocessed data is then resized to match the voxel size of 3D StarDist training data. **(2)** Organoid segmentation was performed using one of the channels or the channels mean, it consists of steps as follows: Enhance Local Contrast (CLAHE), Gaussian blur, Otsu threshold, Morphological operations, Keep largest object. **(3)** Nuclei are segmented with an AI Stardist pretrained network. **(4)** The segmented organoid mask is used for cleaning, i.e., debris with centroids outside the organoid are removed and border nuclei are dilated to include them in the organoids. **(5)** Cell segmentation was performed using seeded watershed based on nuclei segmentation and cell channel (nuclei are considered as seeds). The expansion of the watershed is limited to the organoid volume. The distance between a nucleus border and the corresponding cell border is constraint to a maximum value (i.e 14 µm). The label mask of cells is filtered with a 3D median filter.

**Supplementary Figure 2:**
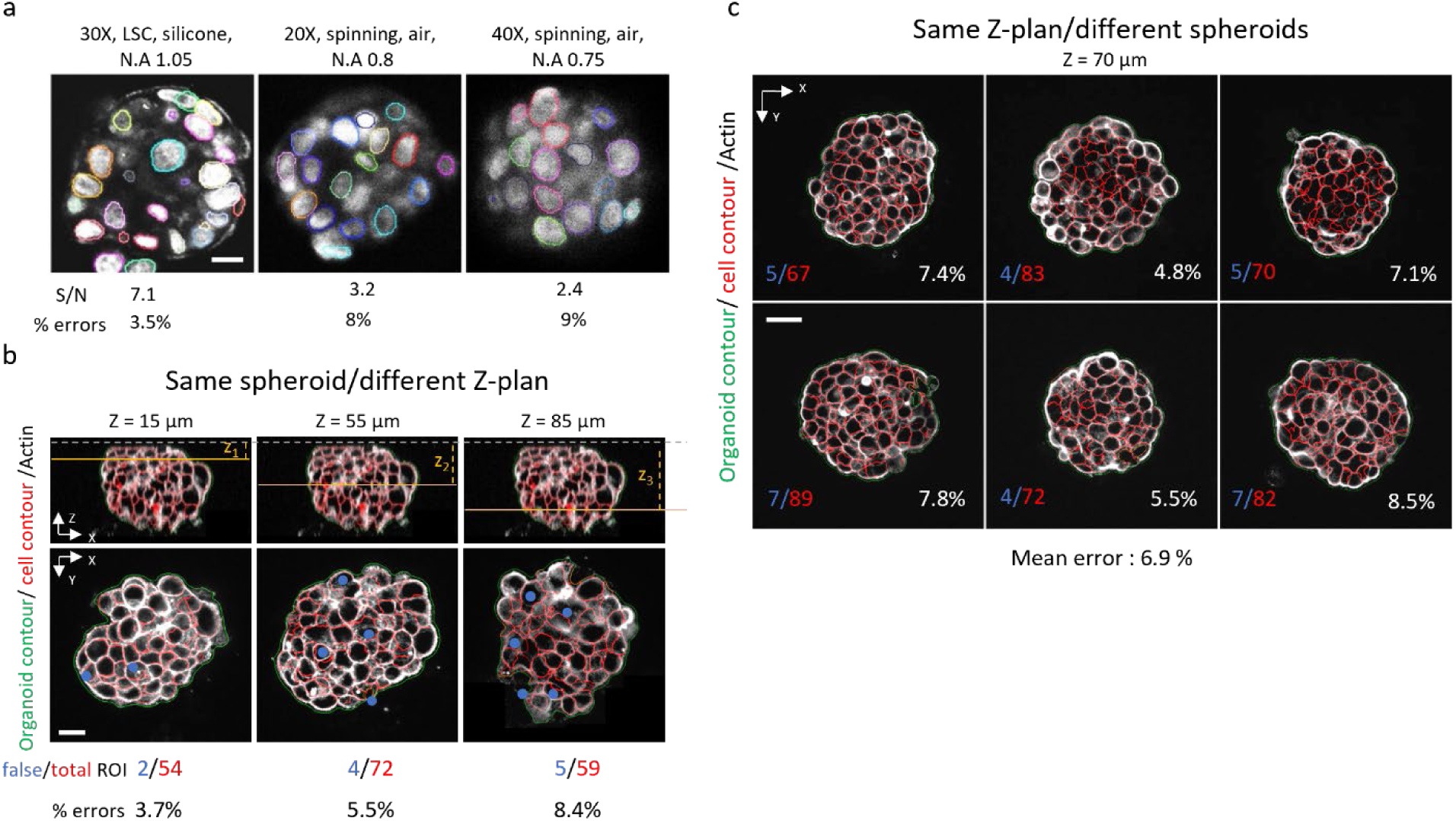
Quality assessment of nuclei and cell segmentation. **a**: Representative segmentation results of nuclei acquired using 3 different configuration and setup. From left to right: Laser Scanning Confocal (LSC) FV3000 Olympus with a 30X, silicone immersion objective, N.A = 1.05. Spinning disk confocal BC43 Andor with a 20X, air objective, N.A=0.8. Spinning disk confocal BC43 Andor with a 40X, air objective, N.A=0.75. Signal-to-noise ratio (S/N) and percentage of missing/bad segmentation (% errors) are provided for each image (see online method for details on % errors calculation). **b**: Representative segmentation results of cells, based on Actin staining, of same spheroid at 3 different z-plans with examples of bad segmentation (‘false’ ROI = blue dots) and percentage of mis-segmentation (% errors) are provided for each image (see online method for details on % errors calculation). **c**: Similar as **b** but from 6 different spheroids at the same depth (70 µm) with corresponding mean of percentage errors.

**Supplementary Figure 3:**
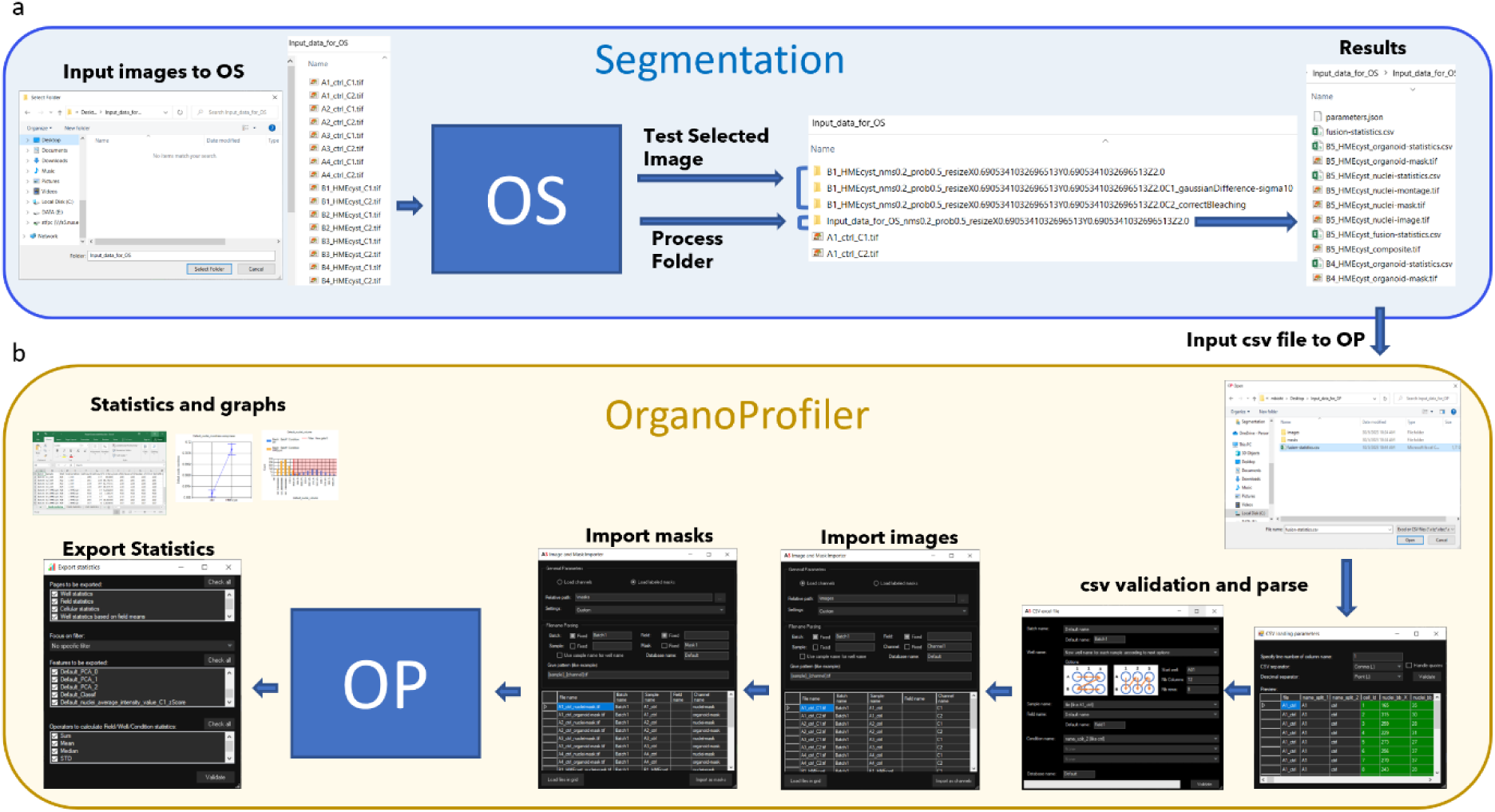
Data import and export. **a:** Segmentation pipeline and files organization, user can test segmentation on selected image or process all images of an input folder. When “Process Folder”, the results for all images were output in one folder. **b:** OrganoProfiler. The “Process Folder” results (.csv) from segmentation can be imported into OrganoProfiler for data mining. Data import starts from csv file validation and parsing, followed by images and masks import. Various statistics and graphs can be exported from OrganoProfiler.

**Supplementary Figure 4:**
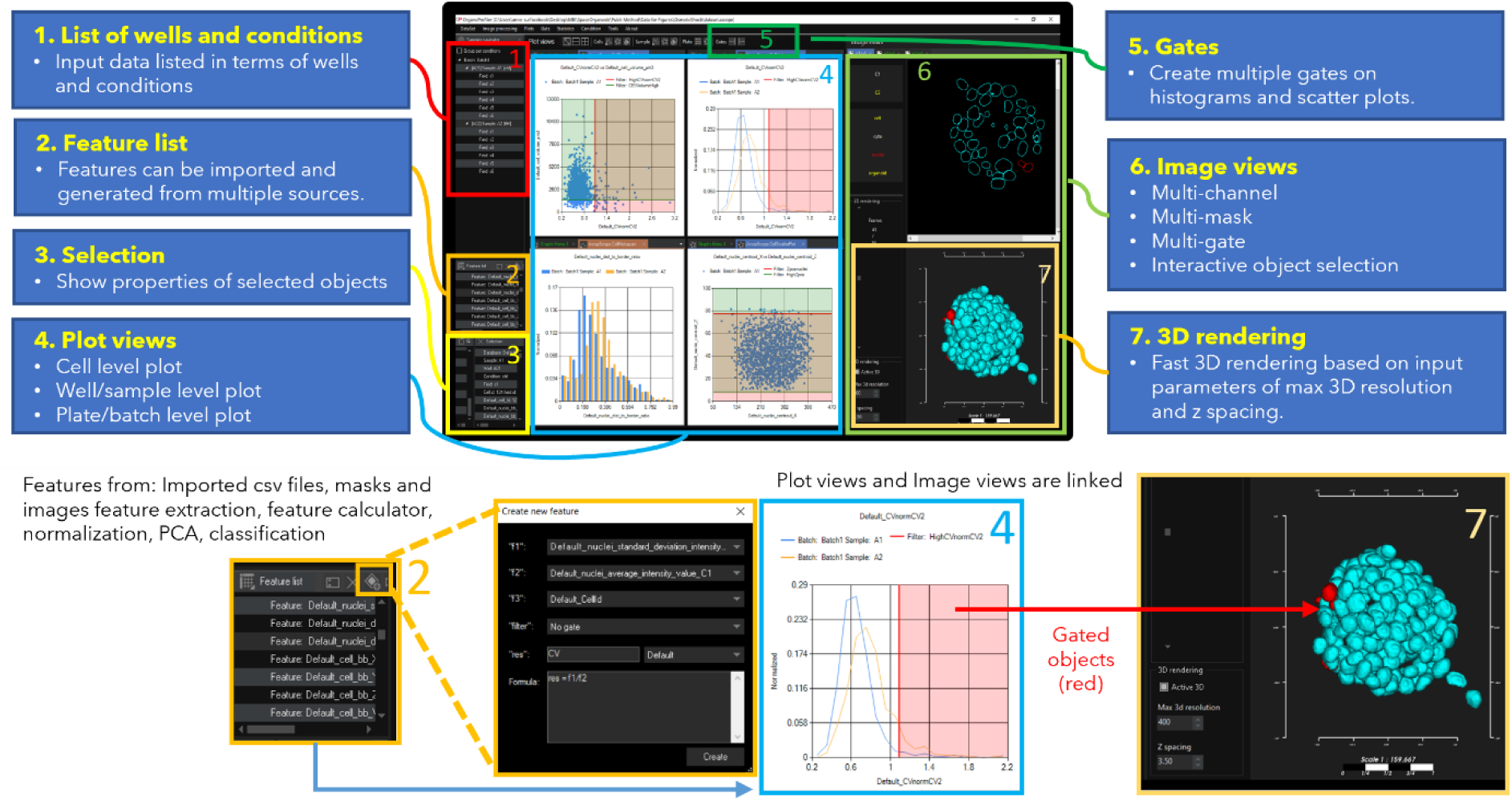
Example of interface to perform data mining. **(1)** List of wells and conditions: Input data listed in terms of wells and conditions. **(2)** Feature list: Features can be imported and generated from multiple sources, e.g., imported csv files, extraction from masks and images, feature calculator, normalization, Principal Component Analysis (PCA), classification (Sport Vector Machine, Decision Tree, Random Forest Algorithms) etc. Feature calculator for creating new feature is shown in the zoom-in view. **(3)** Selection: Selection window shows properties of selected object. Properties shown include sample, well, condition, field and all measurements. **(4)** Plot views: Multiple types of graphs can be created, e.g., cell level plot (Histogram, scatter plot, bubble plot), well/sample plot (Bar plot, scatter plot, bubble plot), plate/batch level plot (Heat map, batch vs. batch scatter plot). **(5)** Gates: Gates can be created on histograms and scatter plots, instant feedback is shown in image views while adjusting gating threshold. **(6)** Image views: Image views show channels, masks and gates. Object can be selected interactively in 2D or 3D or plot view. **(7)** 3D rendering: Fast 3D rendering based on input parameters of max 3D resolution and z spacing.

**Supplementary Figure 5:**
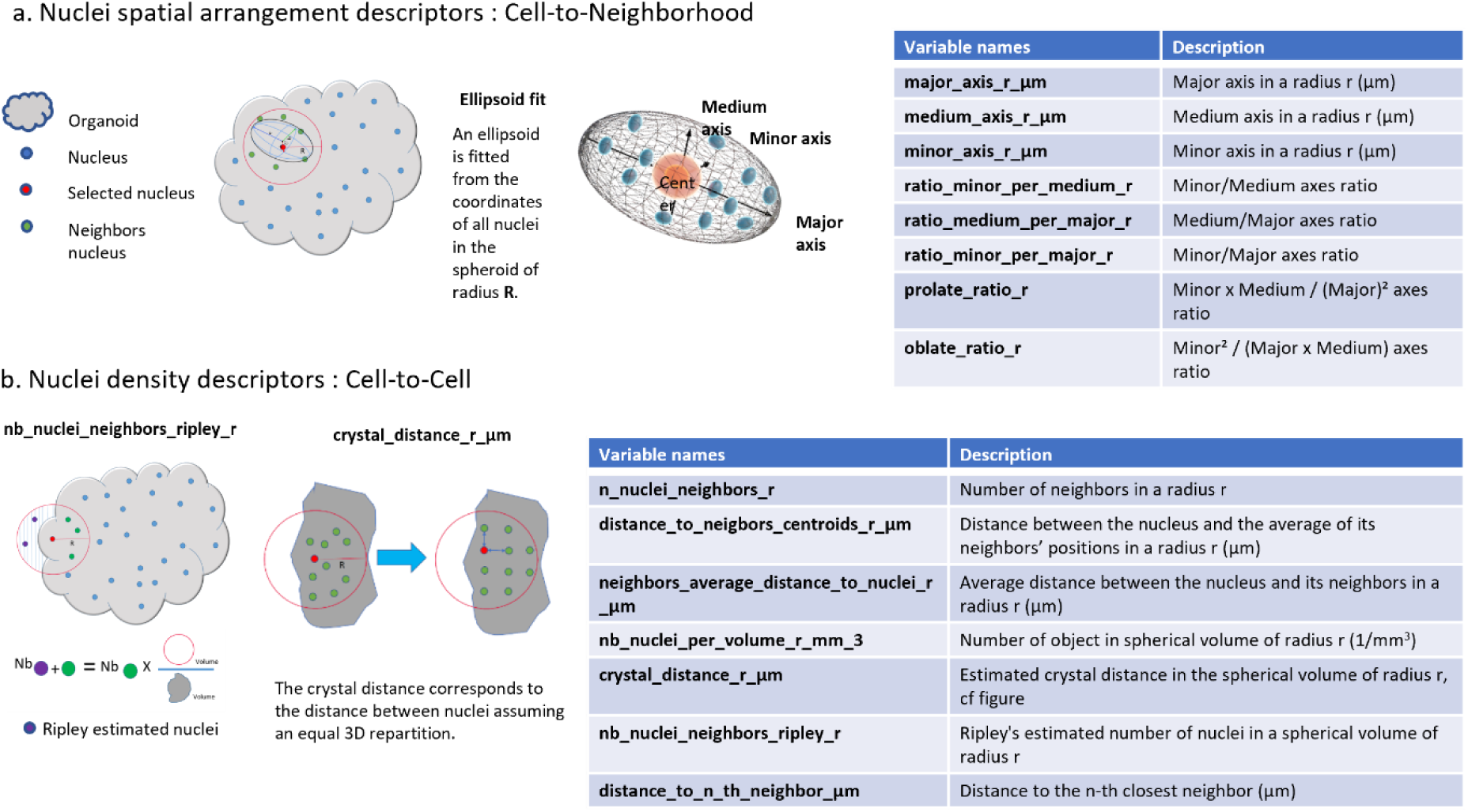
Summary of 3D topological descriptors. **a**. Nuclei spatial arrangement descriptors: Cell-to-Neighborhood: An ellipsoid fitted from all nuclei coordinates in a spheroid of radius r (µm) provides various descriptors (listed in the table). **b**. Nuclei density descriptors: Cell-to-Cell: Illustrations showing Ripley nuclei and crystal distance definition. Ripley nuclei can be estimated by multiplying the number of nuclei inside the sphere of radius r µm with the ratio of sphere volume to organoid volume inside the sphere. Crystal distance is defined as the grid spacing assuming equal 3D repartition of nuclei in the spheroid. The full list of nuclei density descriptors is shown in the corresponding table.

**Supplementary Figure 6:**
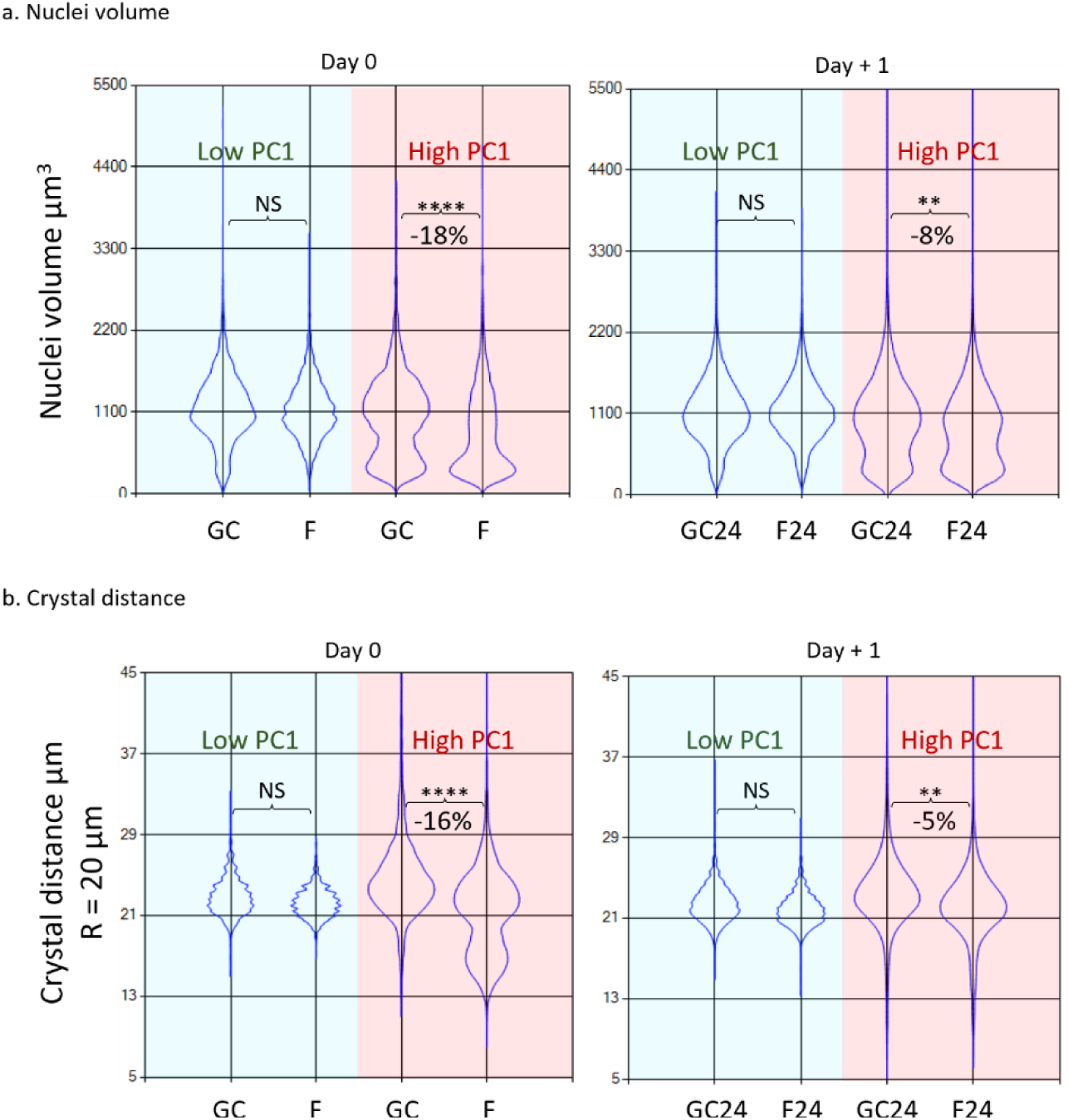
Violin plots with unsupervised clustering of PCA classification. **a**. Differences in cell nuclei volume among conditions (Day 0: GC and F, Day+1: GC24 and F24) and with PC1 classification (Red: high PC1 value, Green: low PC1 value) (****: P<0.0001, **: P<0.01). **b**. Similar as **a**. but for differences in crystal distance within a radius (R) of 20 µm.

